# Methylmalonic acid impairs cell respiration and glutamate uptake in C6 rat glioma cells

**DOI:** 10.1101/2021.08.26.457835

**Authors:** Renata T. Costa, Marcella B. Santos, Carlos Alberto-Silva, Daniel C. Carrettiero, César A.J. Ribeiro

## Abstract

Methylmalonic acidemia is an organic acidemia caused by deficient activity of L-methylmalonyl-CoA mutase or its cofactor cyanocobalamin and it is biochemically characterized by an accumulation of methylmalonic acid (MMA) in tissue and body fluids of patients. The main clinical manifestations of this disease are neurological and observable symptoms during metabolic decompensation are encephalopathy, cerebral atrophy, coma, and seizures, which commonly appear in newborns. This study aimed to investigate the toxic effects of MMA in a glial cell line presenting astrocytic features. Astroglial C6 cells were exposed to MMA (0.1-10mM) for 24 or 48 hours and cell viability, glucose consumption and oxygen consumption rate, as well as glutamate uptake and ATP content were analyzed. The possible preventive effects of bezafibrate were also evaluated. MMA significantly reduced cell viability after 48-hour period and increased glucose consumption during the same period of incubation. Regarding the energy homeostasis, MMA significantly reduced respiratory parameters of cells after 48-hour exposition, indicating that cell metabolism is compromised at resting and reserve capacity state, which might influence the cell capacity to meet energetic demands. Glutamate uptake and ATP content were also compromised after exposition to MMA, which can be influenced energy metabolism impairment, affecting the functionality of the astroglial cells. Our findings suggest that these effects could be involved in the pathophysiology of neurological dysfunction of this disease.

## INTRODUCTION

Methylmalonic acidemia is a severe inherited metabolic disorder of intermediary metabolism caused by defective activity of the enzyme L-methylmalonyl-CoA mutase (MCM) or by defects in the synthesis of its cofactor adenosylcobalamine. MCM catalyzes the conversion of L-methylmalonyl-CoA to succinyl-CoA, a key step in the catabolism of branched-chain amino acids, pyrimidines, odd-chain fatty acids and cholesterol side-chains (Deodato et al. 2006; Van Gosen 2008). The defect in MCM activity leads to the accumulation of methylmalonic acid (MMA), among other toxic compounds, in tissue and body fluids (Fenton et al. 2001; Zwickler et al. 2012; Haijes et al. 2019b). This disease has an estimated incidence of one in 50000 individuals (Baumgartner et al. 2014).

The biochemical profile of affected patients is characterized by metabolic acidosis, hyperammonemia, hyperglycinemia and hypoglycemia (Fenton et al. 2001; Haijes et al. 2019a). Clinical manifestations are highly variable, depending on the severity of the enzyme defect, and include mental retardation, cerebral atrophy, coma and seizures and are frequently observed in the first months of life (Horster et al. 2007). Patients who survive to decompensation episodes usually present a variable range of psychomotor and cognitive delays, demyelination as well as cortical and basal ganglia damage (van der Meer et al. 1994; de Baulny et al. 2005; O’Shea et al. 2012).

Although various investigations have suggested the toxicity of accumulating metabolites, the pathophysiological mechanisms of the brain damage are not completely understood. Studies demonstrated that MMA is toxic to rodent central nervous system (CNS), compromising mitochondrial bioenergetics (Wajner, Coelho 1997; Brusque et al. 2002; Okun et al. 2002; Maciel et al. 2004; Kowaltowski et al. 2006; Pettenuzzo et al. 2006), redox homeostasis (Fontella et al. 2000; Pettenuzzo et al. 2003; Furian et al. 2007) and glutamatergic system (Kolker et al. 2000; de Mello et al. 1996). Energetic dysfunction and oxidative stress were also observed in patients and in knockout models of methylmalonic acidemia (Treacy et al. 1996; Richard et al. 2007; Richard et al. 2009; Chandler et al. 2009; de Keyzer et al. 2009). It was also showed that MMA causes apoptotic and necrotic death in neuronal cultures (Okun et al. 2002; Kowaltowski et al. 2006; McLaughlin et al. 1998), which can be prevented with glutamatergic antagonists, suggesting the involvement of excitotoxic mechanisms (Okun et al. 2002; McLaughlin et al. 1998). Besides the studies showing the toxicity of MMA to neuronal cultures, the effects of this organic acid to astrocyte cells remain to be elucidated.

In this context, astrocytes, the most abundant glial cells in many parts of the CNS (Pakkenberg, Gundersen 1988), interact with neurons, blood vessels, and many structures of this system (Abbott et al. 2006; Cheslow, Alvarez 2016). Since astrocytes are the only cells in the CNS that store and process glycogen, they contribute to the supply of glucose and energy metabolites to neurons, as well as neurotrophins and antioxidant defenses (Maragakis, Rothstein 2006; Souza et al. 2019). They also have many regulatory functions in brain homeostasis, controlling uptake of neurotransmitters (Mahmoud et al. 2019; Danbolt 2001), that is vital for protecting neurons against excitotoxicity. In addition, astrocyte damage due to trauma, ischemia and genetic or metabolic disorders can dramatically affect normal maturation, thereby increasing the probability of impaired brain development (De Keyser et al. 2008; Sidoryk-Wegrzynowicz et al. 2011).

Therefore, in the present study we investigated the effects of MMA on C6 astroglial cells, which are widely used as an astrocyte-like cell line to evaluate astrocytic parameters (Galland et al. 2019; Tramontina et al. 2012; Morioka et al. 2016). We determined the time-course effects of MMA on cell viability, glucose consumption and lactate release, oxygen consumption, superoxide production, reduced glutathione concentrations and malondialdehyde levels, as well as glutamate uptake. The protective effects of bezafibrate, a synthetic agonist of peroxisome proliferator-activated receptor (PPAR), on MMA-elicited effects were also evaluated.

## MATERIAL AND METHODS

### Reagents

Fetal bovine serum (FBS), penicillin/streptomycin and trypsin were purchased from Gibco (Carlsbad, CA, USA). C6 astroglial cells were provided by Prof. André Quincozes-Santos, Dept. of Biochemistry, UFRGS, Brazil. Methylmalonic acid, Dulbecco’s modified Eagle’s medium (DMEM, containing 5.5 mM glucose), methyl-thiazolyl diphenyl tetrazolium bromide (MTT), as well as all other chemicals of analytical grade were obtained from Sigma (St Louis, MO, USA). L-[2,3-^3^H]-glutamate, purchased from PerkinElmer (Boston, MA, USA).

### Cell Culture

Cells were seeded in 75 cm^2^ flasks and maintained at 37°C in an humified 5% CO_2_ and 95% air atmosphere in complete medium (DMEM, 15 mM HEPES, 14.3 mM NaHCO_3_, and antibiotics (100 IU/ml penicillin and 100 μg/ml streptomycin)) supplemented with 10% fetal bovine serum (FBS). Due to the exponential growth of this cell line (doubling times between 24 and 36 h), the cells were seeded every two or three days into new culture flasks with fresh complete medium.

### Experimental design – Methylmalonic acid exposition to cells

Confluent cells were detached from the culture flasks using 0.05% trypsin/EDTA, collected by centrifugation for 5 min (400 x g, 23°C), suspended in complete medium supplemented with 10% FBS and seeded (9 × 10^3^ cells/cm^2^) in 96-, 12-, 6-well plates or 25 cm^2^ flasks, according to each evaluated parameter. After cells reached 80% of confluence, the culture medium was removed and cells incubated in the presence of 0.1 to 10 mM MMA, prepared in DMEM with 5% FBS (pH was adjusted to 7.4 with NaOH) for 24 or 48 hours at 37°C in a 95% air/5% CO_2_ incubator. Control cells were incubated for the same time periods in DMEM supplemented with 5% FBS. After the end of the incubation periods, the culture media was collected to evaluate of glucose consumption and lactate release and the cells were used for the various biochemical analyses. To evaluate the effects of the PPAR agonist bezafibrate (Beza), cells were treated with Beza (200 nM to 500 μM) for 48 hours. The possible protective effect of Beza was determined by co-incubating Beza (200 nM) for 48 hours with MMA (1.0 and 5.0 mM).

### Metabolic activity and cell viability

Metabolic activity was accessed in 96-well plates by the MTT reduction assay, in which formazan is produced by mitochondrial dehydrogenases of functional cells. At the end of the incubation periods, the culture media was changed by fresh culture media (devoid of MMA) containing 0.5 mg/ml of MTT. After 1 h of incubation, the medium from each well was gently removed by aspiration, and 100 μL isopropanol was added to each well followed by incubation and shaking for 5 min to solubilize the formazan product and absorbance measured at 570nm. Data were expressed as percentage of control.

For determination of viability at the end of the incubation periods, cells grown in 12-well plates were detached with trypsin/EDTA, collected by centrifugation and cell viability was determined by flow cytometry with Muse^®^ Count &Viability Assay Kit (Millipore), following the manufacturer’s instructions. Results were expressed as percentage of viable and dead cells.

### Glucose Consumption and Lactate Release

Glucose and lactate concentrations were determined in the culture medium collected after 24 or 48 hours of exposition of cells to MMA. Glucose consumption was determined by the difference between the concentration of glucose at the end of incubation period from the concentration measured at the start of incubation period. Glucose and lactate concentrations we determined by the glucose oxidase and lactate oxidase methods, respectively, using commercial kits (Labtest, Brazil) and were expressed as nmol glucose / hour or mM lactate in culture media.

### High Resolution Respirometry

The oxygen consumption rate (OCR) was measured by high-resolution respirometry using Oroboros O2K Oxygraph (OROBOROS Instruments, Innsbruck, Austria) in 2 mL chamber at a stirrer speed of 750 rpm. Preliminary experiments were carried out to achieve the best experimental conditions to determine OCR in C6 cells. These conditions included the optimal number of cells per analysis and the concentrations of uncoupler and inhibitors needed to achieve maximal uncoupling and OCR inhibition. Manual titration of inhibitors and uncoupler was performed using Hamilton syringes (Hamilton Company, Reno, NV, USA). Before each experiment, the oxygen concentration in the medium was equilibrated with air in the respirometer chambers at 37°C until a stable signal was obtained at an oxygen concentration of approximately 190 μM.

After the exposition to MMA for 24 or 48 hours, cells were detached from 25 cm^2^ flasks using 0.05% trypsin/EDTA, collected by centrifugation for 5 min (400 x g, 23°C), suspended in 300 μL of DMEM supplemented with 5% FBS and counted in an hematocytometer. OCR was then measured by incubating the cells (1.0x 10^6^) at 37°C in a 2 mL chamber containing DMEM supplemented with 5% FBS.

Respiratory states were determined by substrate-uncoupler-inhibitor titration (SUIT) protocol according to Pesta and Gnaiger (2012). In brief, basal OCR, reflecting the aerobic metabolic activity under cellular routine conditions, was determined by monitoring OCR for 5 min after a stable signal was achieved. Inhibitors and uncouplers were then added to the chambers through the injection port of the stoppers using Hamilton syringes. Leak-related OCR was measured after addition of oligomycin A (0.5 μM, final concentration) and indicates the oxygen consumption to compensate proton leak after inhibition of ATP synthase. The addition of the uncoupler CCCP (4.5 μM, final concentration) allows the determination of the maximal OCR, reflecting the maximal capacity of the electron transfer system. Residual oxygen consumption (ROX), related to non-mitochondrial oxygen consumption enzymes, was determined after the addition of the inhibitors rotenone and antimycin A (0.5 μM each, final concentrations).

To investigate the mitochondrial respiration, the following respiratory parameters were calculated: (I) Routine respiration (basal oxygen consumption employed to supply cellular regular metabolism), (II) proton leak respiration (oxygen consumption related to proton flux across the mitochondrial inner membrane), (III) ATP-linked respiration (amount of oxygen consumed by cells to produce ATP, determined by the rate of respiration that can be inhibited by oligomycin), (IV) Maximal respiration (oxygen consumption after mitochondrial uncoupling, related to maximal electron flux in the electron transfer chain), (V) reserve respiratory capacity (the difference between maximal and routine respiration) and (VI) non-mitochondrial respiration (ROX, related to oxygen consumption after inhibition of complexes I and III of the electron transfer chain).). Data recording was performed using the DataLab software 7.04 (OROBOROS Instruments). Oxygen flux in all the respiratory parameters was corrected for non-mitochondrial respiration (ROX) and expressed as pmol O_2_ / s / 10^6^ cells.

### Oxidative Stress Parameters

The induction of oxidative stress was evaluated after exposition of cells to MMA for 48 hours. For the determination of superoxide production, cells grown in 12-well plates were detached with trypsin/EDTA, collected by centrifugation and superoxide anion production was determined by flow cytometry with Muse^®^ Oxidative Stress Assay Kit (Millipore), following the manufacturer’s instructions. Results were expressed as fluorescence arbitrary units of dihydroethidine.

For determination of malondialdehyde and GSH concentrations, cells grown in 6-well plates were scraped 20 mM sodium phosphate buffer, pH 7.4, containing 140 mM KCl, and then were centrifuged at 750 g (10 min, 4°C) to discard nuclei and cell debris, and supernatant, a suspension of preserved organelles, including mitochondria, was used (Olivera-Bravo et al. 2015).

MDA levels were determined according to Yagi (1998) with slight modifications. Briefly, 200 μl of 10% trichloroacetic acid and 300 μl of 0.67% thiobarbituric acid in 7.1% sodium sulfate were added to 100 μl of cell supernatants and incubated for 2 h in a boiling water bath. The mixture was allowed to cool in running tap water for 5 min. The resulting pink-stained complex was extracted with 300 μl of butanol. Fluorescence of the organic phase was read with 515 nm excitation and 553 nm emission wavelengths. MDA levels, determined in triplicate for each experimental condition, were calculated as nmol MDA/mg protein using a calibration curve determined with 1,1,3,3-tetramethoxypropane.

GSH concentrations were measured according to Browne and Armstrong (1998) with minor modifications. Supernatants (0.3–0.5 mg of protein/mL) were first deproteinized with metaphosphoric acid, centrifuged at 7000g for 10 min and immediately used for GSH quantification. One hundred and eighty-five microliters of 100 mM sodium phosphate buffer, pH 8.0, containing 5 mM ethylenediaminetetraacetic acid, and 15 μl of o-phthaldialdehyde (1 mg/mL) were added to 30 μl of supernatant previously deproteinized. This mixture was incubated at RT in a dark room for 15 min. Fluorescence was measured using excitation and emission wavelengths of 350 and 420 nm, respectively. The calibration curve was prepared with standard GSH (0.001–1 mM) and the concentrations, determined in triplicate for each experimental condition, and referred as nmol GSH/mg protein).

### Glutamate uptake

Glutamate uptake was performed as previously described (Galland et al. 2019). After 48 hours exposition to MMA, the medium was changed and C6 cells were incubated at 37 °C in HBSS buffer containing the following components (in mM): 137 NaCl, 5.36 KCl, 1.26 CaCl_2_, 0.41 MgSO_4_, 0.49 MgCl_2_, 0.63 Na_2_HPO_4_, 0.44 KH_2_PO_4_, 4.17 NaHCO_3_ and 5.6 glucose, pH adjusted to 7.4. The assay was started by the addition of 0.1 mM L-glutamate and 0.33 μCi/ml l-[2,3-^3^H] glutamate. The incubation was stopped after 10 min by removing the medium and rinsing twice with ice-cold HBSS. The cells were then lysed in a solution containing 0.5 M NaOH. Incorporated radioactivity was measured in a scintillation counter. Sodium-independent uptake was determined using N-methyl-D-glucamine instead of sodium chloride. Sodium-dependent glutamate uptake, considered specific uptake, was obtained by subtracting the sodium-independent uptake from the total uptake. Glutamate uptake was calculated as nmol glutamate /min /mg protein and was expressed as percentage of control.

#### ATP Content

For determination of ATP content, cells were grown in 6-well plates. After exposition to MMA for 48 hours, the medium was removed, cells were rapidly washed with cold PBS to remove medium and 200 μL cold 0.6 M perchloric acid was added to each well. Plates were kept on ice for 15 minutes for metabolic extraction. Wells were scraped, the lysate transferred to an Eppendorf tube and centrifuged for 10 mins at 20.000 x g to sediment protein content. Supernatants were transferred to new Eppendorf tubes and pH neutralized with 2M K_2_HPO_4_, whereas pellets were solubilized with 0.1 NaOH for further protein content determination. After addition of K_2_HPO_4_, tubes were kept on ice for 15 mins and centrifuged for 10 mins at 20.000 x g to sediment potassium perchlorate crystals. Supernatants were stored at −80°C until ATP content determination, which was carried out by luminescent Promega ENLITEN ATP kit (Promega, USA). Data were calculated as pmol ATP / mg protein and expressed as percentage of control.

### Protein Content

Protein content was measured according to Bradford (1976), using bovine serum albumin as a standard.

### Statistical Analysis

Data are expressed as mean ± SEM (n=4-6 as described in figure legends). Comparisons between two groups were performed with unpaired *Student t*-tests. Comparisons between multiple groups were performed with One-way ANOVA with Tukey’s post hoc analyses for comparisons between control and treatment groups. A *p* value of less than 0.05 was considered significant. All the statistical analyses were performed with Prisma Software (version 6.07, GraphPad Software Inc, San Diego, CA).

## RESULTS

### Methylmalonic acid compromises C6 astroglial cells mitochondrial function but does not cause cell death

As a first experimental approach, we investigated the effects of MMA to astroglial C6 cells by measuring MTT reduction. Formazan production was not altered after 24 hours MMA exposition (Figure 1A). MMA, in concentrations ranging from 1.0 to 10 mM, significantly decreased MTT reduction in 48-hour period (30%, *p* <0.01, Figure 1B), suggesting that this metabolite compromises mitochondrial function in a time-dependent way. However, when cell viability was determined by flow cytometry, no changes were observed in the percentage of viable and dead cells exposed to MMA for 24 or 48 hours (Figures 1C and 1D).

**Fig 1.**
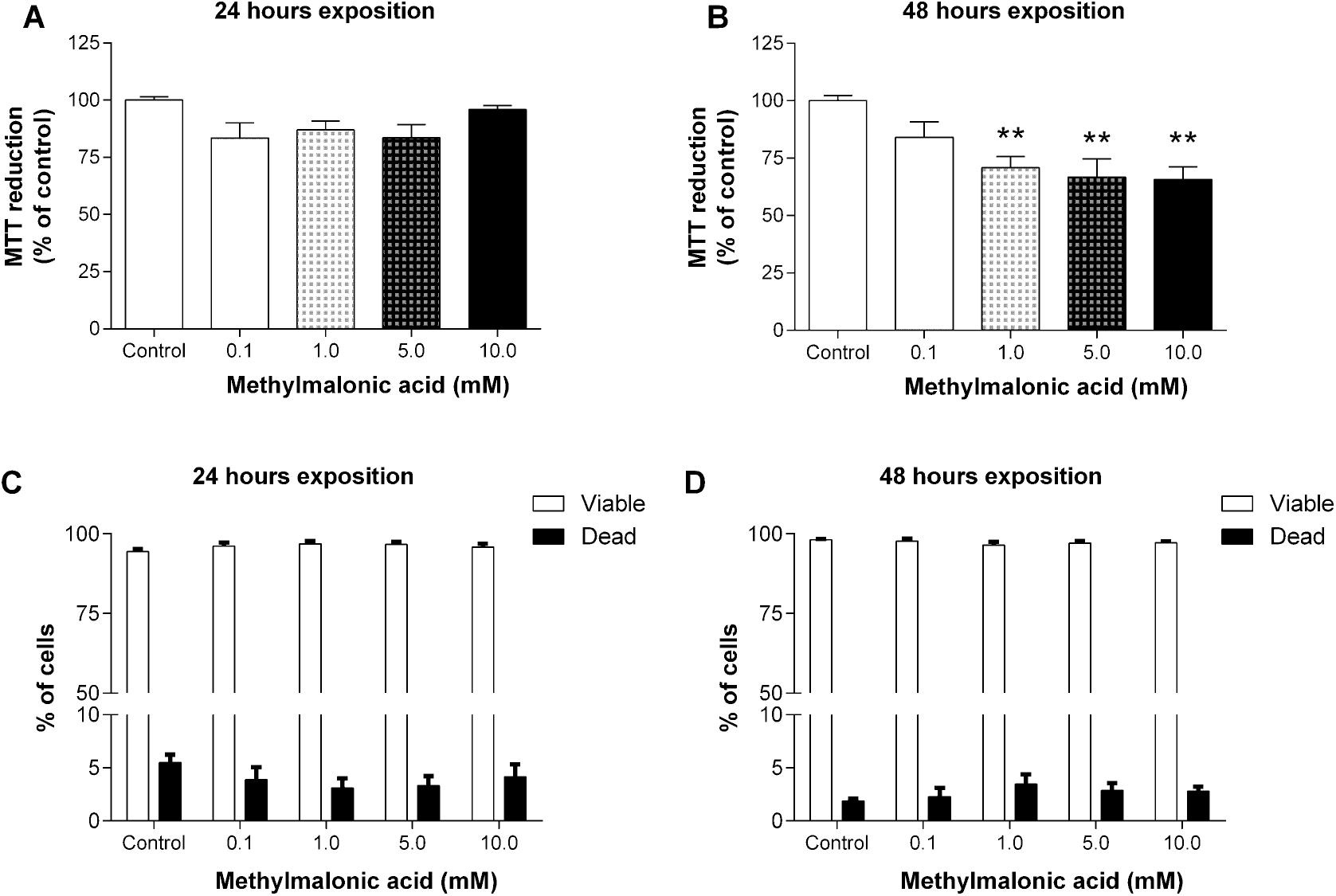
Effects of methylmalonic acid (MMA) on metabolic activity and cell viability. Metabolic activity was measured by MTT reduction assay whereas cell viability was measured by flow cytometry after exposition of C6 glioma cells to MMA for 24 hours (panels A and C) or 48 hours (panels B and D). Statistical analysis was performed with one-way ANOVA and multiple comparisons with Tukey’s post-hoc test. Data are presented as percentage of controls, and expressed as mean ± SEM of 4-5 independent experiments. ** *p* < 0.01 compared to control group.

### MMA increases glucose uptake but does not change lactate release in C6 astroglial cells

The effects of MMA exposition on glucose uptake and lactate release in C6 astroglial cells were also examined. MMA caused an increase of glucose uptake in a time and dose-dependent fashion, since no alterations were observed at 24-hour exposition (Figure 2A), but 48-hour exposure significantly increased glucose uptake (35-50%, *p* < 0.01, Figure 2B). A non-significant increase (20%) in lactate concentration in culture media was only observed in cells exposed to 10 mM MMA (Figure 2C).

**Fig 2.**
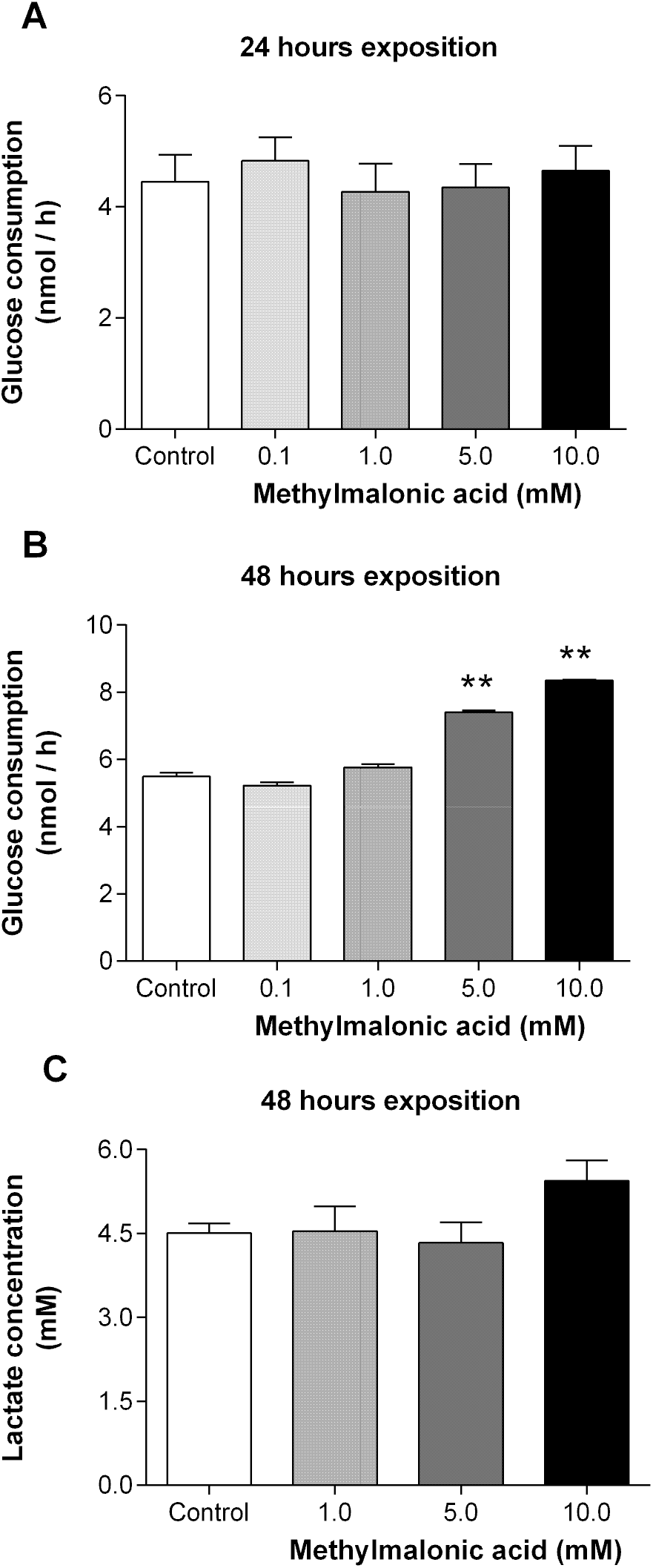
Glucose consumption and lactate release by C6 glioma cells exposed to methylmalonic acid (MMA). The effects of MMA on glucose consumption were evaluated after 24-hour (A) and 48-hour (B) exposition. Lactate release was evaluated after 48-hour (C) exposition to MMA. Statistical analysis was performed with one-way ANOVA and multiple comparisons with Tukey’s post-hoc test. Data are presented as nmol glucose consumed / hour or mM lactate released, and expressed as mean ± SEM of four independent experiments. ** *p* < 0.01 compared to control group.

### Methylmalonic acid compromises respiration in C6 astroglial cells

Considering that a decrease in MTT reduction and an increase in glucose consumption may suggest alterations in mitochondrial functioning, the next step of our study was to determine its effects on C6 cell respiration. No alterations in the respiratory parameters were observed after 24-hour exposition of cells to MMA (data not shown). However, as shown in Figure 3, after 48-hour exposition, MMA significantly reduced Routine Respiration (panel A, 20-45%, *p* < 0.01), ATP-linked Respiration (panel B, 25-50%, *p* < 0.01), Maximal Respiration (panel C, 38-57%, *p* < 0.001) and Reserve Respiratory Capacity (panel D, 50-70%, *p* < 0.01), without altering proton leak and ROX (data not shown).

**Fig 3.**
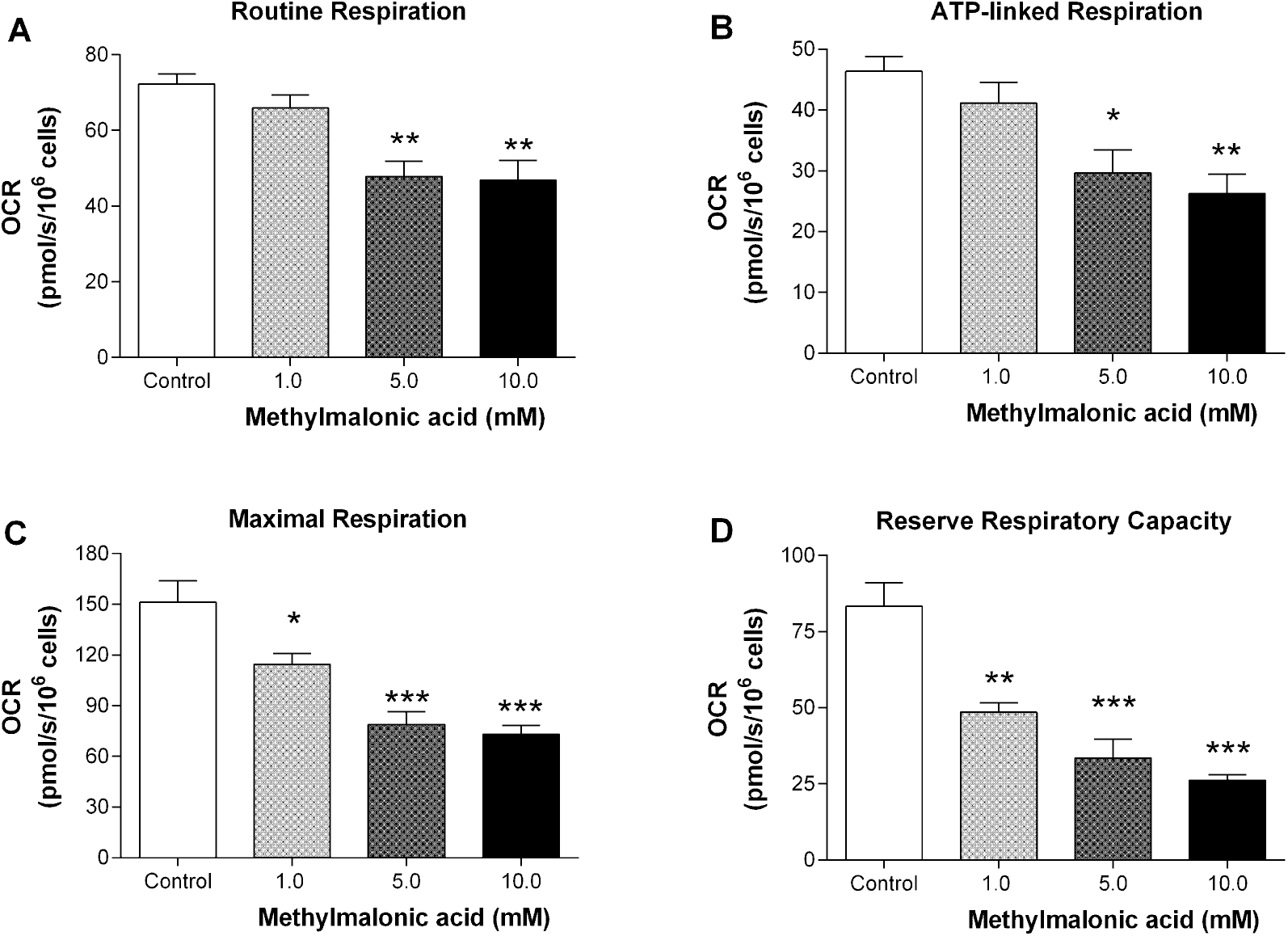
Oxygen consumption rate (OCR) in C6 glioma cells exposed to methylmalonic acid (MMA) for 48 hours. Routine Respiration (A), ATP-linked respiration (B), Maximal respiration (C) and Reserve Respiratory Capacity (D) were determined as described in Material and Methods section. Statistical analysis was performed with one-way ANOVA and multiple comparisons with Tukey’s post-hoc test. Data are represented as pmol consumed O_2_ / s / 10^6^ cells, and expressed as mean ± SEM for of 4-5 independent experiments. * p<0.05, ** p<0,01, *** p<0.001 compared to control group.

### Methylmalonic acid does not alter oxidative stress parameters in C6 astroglial cells

The production of superoxide anions, levels of reduced glutathione and malondialdehyde were not altered in C6 cells exposed to MMA for 48 hours, suggesting that at our experimental scenario, MMA does not induce oxidative stress (data not shown).

### Bezafibrate does not alter the reduction in cell respiration induced by MMA

Aiming to prevent the inhibitory effects of MMA in cell respiration, the effects of bezafibrate, a PPAR agonist with the ability to induces mitochondrial biogenesis (Augustyniak et al. 2019), were evaluated. To determine the lowest dose of bezafibrate that does not alter respiration, C6 cells were exposed to bezafibrate, in concentrations ranging from 200 nM to 500 μM, for 48 hours and the respiratory parameters analyzed. Table 1 shows that bezafibrate reduced all respiratory parameters in a dose-dependent way.

**Table 1:** Respiratory parameters evaluated in C6 glioma cells exposed to increasing concentrations of bezafibrate (Beza) for 48 hours.

|  | Routine Respiration | ATP-Linked Respiration | Maximal Respiration | Spare Respiratory Capacity |
| --- | --- | --- | --- | --- |
| Control | 63.4 ± 3.96 | 46.2 ± 3.21 | 123.2 ± 5.28 | 56.8 ± 4.45 |
| Beza 200 nM | 66.7 ± 3.20 | 43.9 ± 3.23 | 115.9 ± 7.95 | 57.7 ± 2.63 |
| Beza 500 nM | 62.6 ± 3.89 | 41.7 ± 3.53 | 104.0 ± 3.32 | 41.4 ± 4.58 * |
| Beza 15 µM | 57.6 ± 4.33 | 37.9 ± 4.51 | 92.7 ± 2.78 * | 35.1 ± 5.55 * |
| Beza 100 µM | 44.4 ± 2.32 ** | 24.2 ± 2.30 ** | 71.3 ± 6.37 ** | 26.9 ± 4.53 ** |
| Beza 250 µM | 42.4 ± 1.67 ** | 22.2 ± 2.81 ** | 68.4 ± 7.71 ** | 26.0 ± 6.69 ** |
| Beza 500 µM | 35.3 ± 1.15 *** | 15.5 ± 0.65 *** | 51.4 ± 4.25 ** | 16.1 ± 3.19 ** |

Values are presented as mean ± SEM (N = 6) and are expressed as pmol O<sub>2</sub> / s / million cells. \*\* P < 0.01; \*\*\* P < 0.001 compared to control group (One-way ANOVA followed by Tukey's multiple comparisons test).

The possible protective effect of bezafibrate on MMA-induced respiration reduction was studied by co-incubation of MMA with 200 nM bezafibrate, the lowest concentration that did not modify cell respiration. Beza partially attenuated MMA effects on routine respiration (Figure 4A, *p* < 0.05), and completely prevented the effects on ATP-linked respiration (Figure 4B, *p* < 0.05), without modifying the deleterious effects of MMA on maximal respiration (Figure 4C) and on reserve respiratory capacity (Figure 4D).

**Fig 4.**
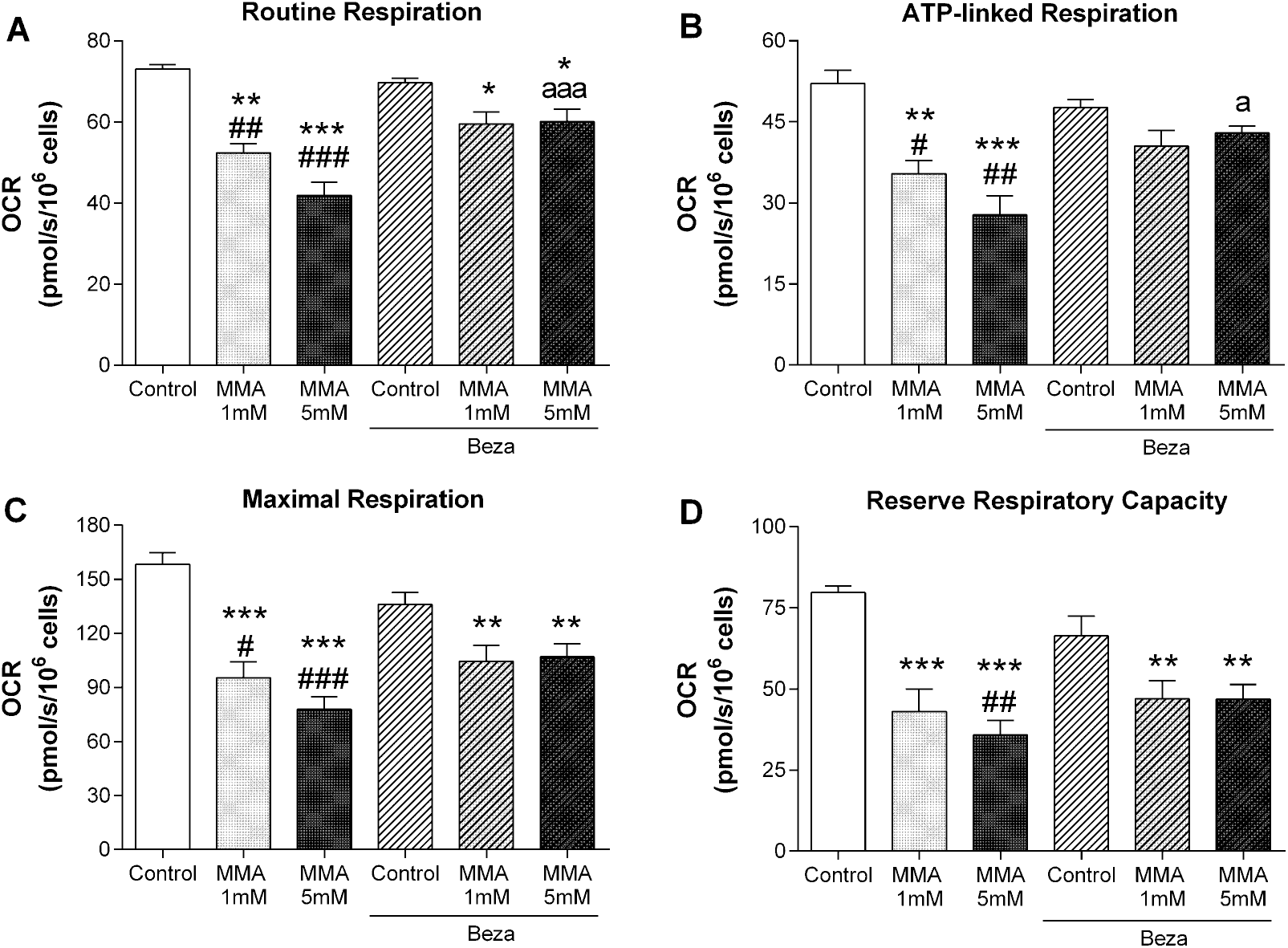
Effects of co-exposition of methylmalonic acid (MMA, 1 and 5 mM) and bezafibrate (Beza, 200 nM) on C6 glioma oxygen consumption rate (OCR). The respiratory parameters Routine Respiration (A), ATP-linked respiration (B), Maximal respiration (C) and Reserve Respiratory Capacity (D) were determined as described in Material and Methods section. Statistical analysis was performed with one-way ANOVA and multiple comparisons with Tukey’s post-hoc test. Data are represented as pmol consumed O_2_ / s / 10^6^ cells, and expressed as mean ± SEM for 4 independent experiments. * p<0.05, ** p<0,01, *** p<0.001 compared to control group; # *p* <0.05, ## *p* < 0,01, ### p<0.001 compared to Beza; a *p* < 0.05; aaa *p* < 0.001 compared to MMA5 (5 mM).

### Methylmalonic acid reduces ATP content and glutamate uptake in C6 astroglial cells

Since neurotransmitter uptake is one of the major functions of astroglial cells (Mahmoud et al. 2019; Danbolt 2001), and this is an energy-dependent process (Gerkau et al. 2017), our next step was to determine the effects of MMA on glutamate uptake and ATP content in C6 astroglial cells. It can be observed in figure 5 that exposition of cells to MMA for 48 hours caused a significant reduction in both glutamate uptake (panel A, up to 15-25%, *p* < 0.01) and ATP levels (panel B, up to 20%, *p* < 0.05), suggesting that this organic acid could compromise the clearance of neurotransmitter from synaptic cleft, which can lead to neurotoxicity.

**Fig 5.**
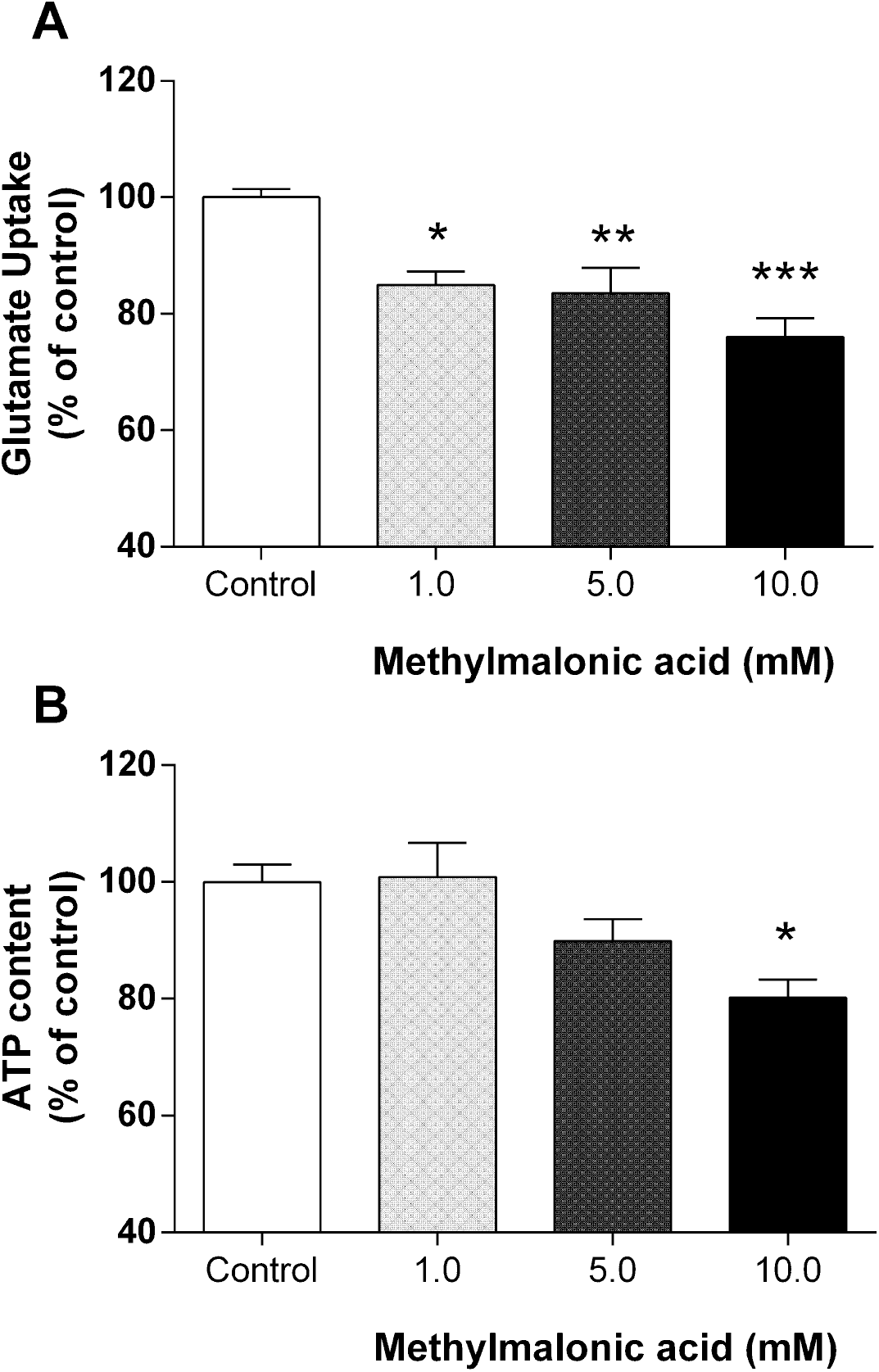
Glutamate uptake and ATP content in C6 glioma cells exposed to methylmalonic acid (MMA) for 48 hours. Statistical analysis was performed with one-way ANOVA and multiple comparisons with Tukey’s post-hoc test. Data are presented as percentage of controls, and expressed as mean ± SEM for 5-6 independent experiments. * *p* < 0.01, ** *p* < 0.01, *** *p* < 0.001 compared to control group.

## DISCUSSION

Astrocytes are the major glial cell type in the central nervous system, participating in a variety of essential physiological processes in the healthy brain, such as in supplying neuronal energy metabolism, controlling ionic balancing and clearance of neurotransmitters (Mahmoud et al. 2019; Souza et al. 2019). Moreover, astrocytes have also been implicated in several neurological disorders (Sidoryk-Wegrzynowicz et al. 2011; Dossi et al. 2018), including organic acidemias (Olivera-Bravo et al. 2016; Olivera-Bravo et al. 2015; Fernandes et al. 2016).

In this scenario, the present study was designed to evaluate the effect of MMA, the main accumulating metabolite in methylmalonic acidemia, on astrocytes. We used C6 glioma cells that are broadly used as astrocyte-like cellular model, since these cell line, at high passages, express astrocytic markers, such as glutamate transporters, glial fibrillary acidic protein, and glutamine synthetase (Galland et al. 2019). For our knowledge, this is the first report of MMA effects on a cell line presenting astrocytic features.

Our first result showed that MMA compromises astrocytic viability measured by MTT reduction assay, in time- and concentration-dependent fashion, without causing cell death (Figure 1). Reductions in cell viability can be caused by several processes: mitochondrial dysfunction, necrosis, autophagia, senescence and/or apoptosis (Tait et al. 2014). Our data suggest that the reduction in cell viability elicited by MMA is associated with a compromising in mitochondrial metabolism, possibly not related to induction of apoptosis or necrosis in this cell type. Moreover, alterations in mitochondrial metabolism were previously reported in vitro (Brusque et al. 2002; Kolker et al. 2000; Kowaltowski et al. 2006; Maciel et al. 2004) and in animal models of methylmalonic acidemia (Chandler et al. 2009; Manoli et al. 2013).

It was also observed an increase in glucose consumption in cells exposed to MMA for 48 hours (Figure 2B), similarly as previously reported in brain of suckling rats (Wajner et al. 1992). Changes in glucose and lactate concentrations were also observed in samples from affected patients (Fenton et al. 2001). Considering the reduction in metabolic viability demonstrated by MTT reduction, together with the increase in glucose consumption, it is feasible to suggest that MMA compromises mitochondrial functioning, and this rise in glucose consumption can indicate an increase in anaerobic glycolysis. An increase in lactate release should also be expected, since this metabolite is a product of increased anaerobic glycolysis. However, we observed only a slight and non-significant increase in lactate release (Figure 2C), which can be explained by the capacity of MMA to direct inhibit lactate dehydrogenase activity (Saad et al. 2006). Moreover, a previous study showed that C6 glioma cells utilize more glucose and produce less lactate than normal astrocytes (Haghighat, McCandless 1997).

Next, we evaluated the effects MMA exposition on cell respiration, which represents a powerful diagnostic model to elucidate mitochondrial function in health and disease. The measurement of oxygen consumption rate by high-resolution respirometry is one of the chosen methods for the analysis of mitochondrial function and dysfunction in culture cells (Brand, Nicholls 2011; Pesta, Gnaiger 2012). Exposition of cells to MMA for 48 hours compromised respiration, since we found a reduction in routine and ATP-linked respiration at 5 and 10 mM and in maximal respiration and reserve respiratory capacity at the lowest tested concentration (1 mM). Routine and ATP-linked respiration are related to oxygen consumption employed to supply cellular regular metabolism at basal state, being dependent on substrate oxidation, ATP turnover and proton leak (Brand, Nicholls 2011). Maximal respiration reflects the maximal electron flux in the electron transfer chain and reserve respiratory capacity describes the amount of extra ATP that can be produced by mitochondria in response to a rapid increase in energy demand, like during ATP-dependent pump activation to maintain ionic potentials and neurotransmitter uptake (Evinova et al. 2020).

Since mitochondrial dysfunctions are related to increase in oxidative stress (Cadenas, Davies 2000), and it was previously demonstrated oxidative damage in MMA patients and study models (Treacy et al. 1996; Chandler et al. 2009; Fontella et al. 2000), we also measured the effects of MMA on formation of superoxide, GSH and malondialdehyde levels. We did not observe alterations in these parameters after exposition of cells to MMA, indicating that our experimental model does not elicit oxidative stress. In this regard, it was also demonstrated in a human neuroblastoma cell model that exposition to MMA for 6 days does not alter GSH content (Stepien et al. 2017).

In an attempt to prevent the toxic effects of MMA on cell respiration, the next set of experiments was designed to evaluate the role of bezafibrate, an activator of PPAR receptor that enhances the levels of PGC-1α coactivator, leading to mitochondrial biogenesis and thus improving mitochondrial function (Augustyniak et al. 2019). Our first approach was to determine a concentration of bezafibrate that did not cause alterations in cell respiration, since mechanisms underlying mitochondrial biogenesis induced fibrates are poorly understood (Pardo et al. 2011). We evaluate the effects of bezafibrate in concentrations ranging from 200 nM to 500 μM, and observed that bezafibrate compromises cell respiration at concentrations as low as 500 nM (Table 1). These results are in line with previous reports demonstrating that fibrates compromise mitochondrial function (Scatena et al. 2004; Brunmair et al. 2004; Wilk et al. 2015; Scatena et al. 2003).

The possible protective effects of Beza on MMA-induced respiration reduction were then studied by co-incubation of MMA with 200 nM bezafibrate, the lowest concentration that did not alter *per se* cell respiration. Beza prevented partially the effects of MMA on routine respiration and completely on ATP-linked respiration, without modifying other respiratory parameters evaluated (Figure 4). It was demonstrated that Beza improved mitochondrial function in neural cells obtained from induced pluripotent human stem cells (Augustyniak et al. 2019) and in fibroblasts of patients with deficiency in fission machinery (Douiev et al. 2020), and in both studies positive effects of Beza were observed in exposition periods longer than used in our study. These data can suggest that the inability of Beza to completely prevent the inhibitory effects elicited by MMA could be caused by shorter exposition period of our study (48 hours). However, it was also demonstrated that long-term exposition to Beza increased markers of mitochondrial diseases in patients (Steele et al. 2020) and an animal model (Yatsuga, Suomalainen 2012), indicating that longer treatments could exacerbate mitochondrial pathologies.

Astrocytes have a crucial role on neurotransmitter homeostasis (Mahmoud et al. 2019), mainly glutamate, the major excitatory neurotransmitter in the CNS (Fonnum 1984; Danbolt 2001). This process highly dependent on energy, since it operates coupled to Na^+^ gradient, that is ultimately dependent on Na^+^,K^+^-ATPase activity (Kirischuk et al. 2016; Magistretti et al. 1999; Rose et al. 2009). It was also demonstrated that mitochondrial dysfunction elicited by several types of toxins can block Na+-dependent glutamate uptake (Robinson, Jackson 2016), emphasizing the direct link between cellular sodium regulation and energy metabolism in the brain.

Considering that MMA reduced cell respiration, we also evaluated whether MMA could compromise astrocytic functionality by measuring glutamate uptake. We found that MMA decreased glutamate uptake (20-25%) in all tested concentrations (Figure 5A). Since it was previously demonstrated in rat synaptosomes that MMA does not directly alter this parameter (Brusque et al. 2001), our results suggest that MMA compromises glutamate uptake by a secondary mechanism, that could be reduction in mitochondrial function or in Na+,K+-ATPase activity. This seems to be true, since inhibitory effects of MMA on Na+,K+-ATPase activity were previously demonstrated (Wyse et al. 2000). Moreover, we also found that MMA reduced cellular ATP content (Figure 5B), which are in agreement with the reduction observed in ATP-linked respiration (Figure 3B).

More importantly, reserve respiratory capacity, which describes the extra ATP that can be produced by mitochondria in response, for example, to ATP-dependent pump activation to maintain ionic potentials and neurotransmitter uptake (Evinova et al. 2020), was also reduced by all MMA tested concentrations (Figure 3D), reinforcing the relationship between mitochondrial dysfunction and reduced glutamate uptake elicited by MMA in our experimental model. Moreover, there is accumulating evidence that inability of astrocytes to deal with increased glutamate concentrations can lead to neuronal death though excitotoxic mechanisms (Kauppinen, Swanson 2007; Gerkau et al. 2017).

Although we did not demonstrate the inhibitory effects of MMA on a specific enzyme or transport system in this study, it was extensively demonstrated deleterious effects of MMA on these systems (Halperin et al. 1971; Dutra et al. 1993; Brusque et al. 2002; Maciel et al. 2004; Pettenuzzo et al. 2006; Mirandola et al. 2008). Compared to these reports, evaluation of cell respiration has an advantage, since it is a measurement of the integrated system, i.e. substrate (glucose) consumption, its conversion to pyruvate in glycolysis, the oxidation of pyruvate in mitochondria and finally the reoxidation of coenzymes in the electron transfer chain by reducing oxygen to water, coupled to ATP synthesis, bringing a greater physiological relevance to our results.

Regarding our experimental model, the tested concentrations of MMA are similar to patients plasma concentrations, which are around 1 mM, but exceed 5 mM due to kidney failure (Kruszka et al. 2013; Manoli et al. 2013). Although the amount of MMA in brain of patients are not known, cerebrospinal fluid concentrations are similar to those found in blood (Hoffmann et al. 1993), and can reach higher levels due to low efflux of dicarboxylic acids through blood-brain barrier, being trapped into the CNS (Kolker et al. 2006). It should be also pointed out that we observed the deleterious effects of MMA only in cells exposed to this compound for longer periods (48 hours). We suppose this is related with the reduced flux of dicarboxylic acids through membranes (Hassel et al. 2002), and MMA takes more time to penetrate the cell and then perform its deleterious effects. Moreover, astrocyte cells express MCM, and its expression can be induced by propionate and isoleucine (Narasimhan et al. 1996), suggesting that a continuous formation and accumulation of MMA occurs in this cell type.

In summary, the results presented in this study demonstrate, for the first time, that MMA compromises cell respiration and glutamate uptake in a cell line presenting astrocyte features in concentrations similar to those found in affected patients, suggesting that astrocyte dysfunction may contribute to the pathophysiology of neurological dysfunction of methylmalonic acidemia.

## DECLARATIONS

## Acknowledgments

Authors would like to thank Prof. Moacir Wajner, Dept. of Biochemistry, UFRGS, Brazil, and Prof. Fernando A.O. Ribeiro, CMCC, UFABC, Brazil, for providing reagents and technical assistance in assays related to radioactive glutamate uptake and ATP content, respectively.

## Availability of data and material

All data are available upon request.

